# Analysis of Heterogeneous Genomic Samples Using Image Normalization and Machine Learning

**DOI:** 10.1101/642108

**Authors:** Sunitha Basodi, Pelin Icer Baykal, Alex Zelikovsky, Pavel Skums, Yi Pan

## Abstract

**Background:** Analysis of heterogeneous populations such as viral quasispecies is one of the most challenging bioinformatics problems. Although machine learning models are becoming to be widely employed for the analysis of sequencing data associated with such populations, their straightforward application is impeded by multiple challenges associated with technological limitations and biases, difficulty of selection of relevant features and need to compare genomic datastes of different sizes and structures.

**Methods:** We propose a novel preprocessing approach to transform irregular genomic data into normalized image data. Such representation allows to restate the problems of classification and comparison of heterogeneous populations as image classification problems which can be solved using variety of available machine learning tools. We then apply the proposed approach to two important molecular epidemiology problems: inference of viral infection stage and detection of viral transmission clusters and outbreaks using next-generation sequencing data.

**Results:** The infection staging method has been applied to HCV *HVR1* samples collected from 108 recently and 257 chronically infected individuals. The SVM-based image classification approach achieved more than 95% accuracy for both recently and chronically HCV-infected individuals. Clustering has been performed on the data collected from 33 epidemiologically curated outbreaks, yielding more than 97% accuracy.

**Availability:** The developed software is freely available at https://bitbucket.org/adv_bio_coll/chronic_vs_clinic

## Background

Currently, viral epidemics continue to be critical public health issues. Many emerging and long-standing epidemics are associated with small (*~* 10 kilobases long) positive-sense single stranded RNA virus, such Human Immunodeficiency Virus (HIV), Hepatitis C virus (HCV), Zika virus (ZIKV) and dengue virus (DENV). The paramount feature of these viruses is their extremely high mutation rate caused by error-prone replication, which can be as high as 10^−4^ [28], thus leading to several expected mutations per every replication cycle. As a result, RNA viruses exist in infected hosts as highly heterogeneous populations of genomic variants usually referred to as *viral quasispecies*. Intra-host and inter-host evolution of viral quasispecies is a complex phenomenon defined by dynamics of virulence, infectivity, drug resistance, immune escape, transmission rates, behavorial patterns and other phenotypic and epidemiological features, which plays crucial role in disease progression and outcome [1, 5, 7, 26, 30]. Challenges associated with understanding complex quasispecies evolution attracted many researchers in different domains, including virology, epidemiology, population genetics and systems biology.

Analysis of heterogeneous viral populations is one of the most challenging bioinformatics tasks due both to the complexity of the underlying algorithmic problems and features and sheer amount of data [2, 22]. These challenges became especially complicated in the recent decade with the advent of high-throughput sequencing (*HTS*), which has now become a major tool for viral research by allowing to sample viral populations on unprecedented depth [3, 10, 14, 16, 19, 29, 31]. Modern computational virology continues mostly to rely on classical approaches, which includes sequence analysis, phylogenetics/phylodynamics and structural bioinformatics [21, 22]. In the recent years, these approaches started to be complemented with the network analysis [6, 13, 32]. Significant number of computational molecular epidemiology problems could be defined using classification- or clustering-based objective. These problems include inference of transmission clusters, detection of co-infections, therapy outcome prediction, infection staging and other research and medical questions. Such problems could be tackled by powerful methods of machine learning and deep learning. It should be expected that in the near future, in accordance with the general trend in AI and Computer Science research, machine learning and deep learning techniques will be utilized in viral research on a much wider scale. However, currently applications of machine learning and deep learning for viral studies is impeded by multiple challenges, which could be thematically classified as follows:

### Challenges associated with technological limitations

High-throughput sequencing technologies and protocols are prone to errors and biases, which are especially pronounced for viral data. Indeed, frequencies of minor viral variants are often comparable with the level of sequencing noise; however, such variants should not be simply discarded based on some frequency threshold, since often they are the ones responsible for transmissions, immune escape or therapy failure [7, 11, 12, 17, 26, 30]. Presence of sequencing errors introduces the noise to the data and produces outlier viral variants, which negatively affect the quality and accuracy of machine learning classifiers.

Another important problem is the sampling and sequencing bias resulting in the significant irregularities in the number and length of viral sequences for different infected individuals. If classifiers capture this artificial differences as significant associations, it may result in overfitting and decline of accuracy after the thorough cross-validation. Thus, application of machine learning to heterogeneous viral population data should be preceded by a preprocessing step, which should be able to eliminate these irregularities through some sort of normalization. However, selection of appropriate normalization approach is challenging. For instance, if we use text classification techniques for preprocessing, then the current difference in number of sequences across different files will either have to be dealt with truncation or padding. This will either cause data loss (in case of truncation) or introduce irrelevant data (in case of padding). An ideal preprocessing method should not introduce any of such issues.

### Challenges associated with feature selection and feature extraction

Before applying machine learning methods for intra-host viral population classification, genomic data of each population should be mapped into the euclidian space ℝ^*n*^. It is usually done by identifying the numerical features that are relevant to the problem under consideration. They can include various diversity measures [24], population genetics parameters [4], physico-chemical properties [21] and other parameters specifically tailored to particular problems. These features are generally identified in consultation with domain experts, and selection of the most relevant features is daunting and resource-consuming task. The role of feature selection in determining the classification performance is paramount. Selection of limited number of features from certain domains inevitably results in loss of information, while increase of feature space dimensionality increases the risk of overfitting and jeopardizes the algorithm’s scalability.

An ideal feature selection method should be able to capture the entire population structure using a relatively simple and easily constructible data representation. Furthermore, it should use a standard universal data format which has fixed number of features and can be applicable to different problems. Since genomic data is essentially a textual information, it is tempting to utilize well-developed machinery from the text classification domain [18, 20] for the purpose of construction of such representation. Viral populations could be mapped to a euclidian spase using word2vec approaches [23], and classified using various available deep learning models [18, 20]. However, application of text processing approaches to viral research could be impeded by several factors. Since they are based on deep neural network models with large numbers of hyperparameters, it requires large annotated datasets to train these models. However, in molecular epidemiology, the amount of available training data is limited in comparison with the text processing domain. The datasets of several hundred intra-host viral populations analyzed in this paper are typical in this context. Although, word2vec or document embedding methods can be directly employed, it is challenging to train a model to get higher classification performance. Furthermore, since viral haplotypes are unique, the trained model could overfit the data.

### Challenges associated with data comparison

Clustering of intra-host viral populations requires an inter-population distance measure, which takes into account complex population structures. It has been shown that among simple alignment-based population distance measures, the minimal distance between population variants allows to achieve the highest clustering accuracy [9]. However, this measure is sensitive to noise and presence of outliers, and does not take into account the whole population structure. Recently, several simulation-based and network-based distance measures have been proposed [13, 32], which overcome above-mentioned limitations at the cost of lesser scalability. Thus, the universal, accurate and efficiently computable inter-population distance measure, which takes into account the complex population structures still has to be developed.

#### Contribution

Several encoding schemes have been used in the literature for coding biomedical data into numerical data for machine learning [37]. In this work, we propose a novel method which converts genomic data into images which are then used for classification. The new approach allows to utilize in genomic analysis the well-developed machine learning methodology from the domain of image processing. The proposed approach allows to address the aforementioned challenges by providing the data structure for the representation of intra-host population structure which is compact, easily adjustable, robust to technological noise, preserve structural properties of populations and can be used for a variety of classification problems, where machine learning is useful.

We validated our approach by applying image processing techniques to two important molecular epidemiology problems. The first problem is the HCV infection staging, i.e. inference whether a patient is recently or chronically infected using viral sequences sampled by next-generation sequencing (NGS). It is known that in *~* 80% of cases untreated HCV infection turns into a chronic infection leading to severe health problems such as liver cirrhosis and hepatocellular carcinoma (a form of liver cancer). HCV infection often does not manifest any symptoms in its early stages, which impedes the timely diagnostic of disease. Furthermore, currently there are no diagnostic assays to determine the stage of an HCV infection. Therefore, distinguishing recently infected patients from chronically infected patients using non-invasive methods such as analysis of genomic data would be highly important both for personalized therapeutic purposes and for the epidemiological surveillance (for example, for detection of incident HCV cases).

The second problem is detection of outbreaks using NGS data. In molecular epidemiology, it is common to utilize the observation that viral populations from the same outbreak are genetically related. Thus, some measure of genetic relatedness could be used as a predictor for epidemiological relatedness [8, 34, 35]. In other words, this problem could be considered as the problem of clustering of intra-host viral populations. Until recently, most available tools for outbreak investigations analyzed only a single representative sequence per population (usually consensus sequence) [34, 35]. Although several recently published tools allow to take into account entire intra-host populations [13, 32, 36], the problem of comparison and clustering of viral populations still remains challenging [15].

We demonstrate that classification and clustering techniques based on normalized image representations of intra-host viral populations allow to solve these two problems with high accuracy.

## Methods

### Data collection

Intra-host HCV populations sampled by sequencing of a highly heterogeneous genomic region (HVR1) were analyzed. The data [4] used for classification of intra-host HCV populations as recent and chronic consists of 365 NGS samples, including 108 datasets correponding to recently infected hosts and 257 datasets belonging to chronically infected hosts. For clustering and indentification of outbreaks we used the benchmark dataset [8, 13, 32] that consists of HCV intra-host populations collected from 335 infected individuals by Centers for Disease Control and Prevention in 2008–2013. Of these, 142 HCV samples belong to epidemiologically curated outbreaks involving from 2 to 19 patients, while the remaining datasets are epidemiologically isolated cases.

### Sequence Image Normalization

We transform sequencing data into an image by the preprocessing method further referred to as Sequence Image Normalization. We assume that sequences are aligned and ordered by their counts, with sequences of the same counts being sorted lexicographically. Next, each symbol *l* ∈ {*′ A′,′ C′,′ T′,′ G′,′* −′} is associated with a particular color thus transforming the sequence alignment into an image. Finally, the images corresponding to different infected hosts are normalized by transforming them into fixed size images. The colors to represent nucleotides are selected from the set of colors of higher variation in order to simplify identification of discriminative features characterizing particular intra-host populations. Fig.1 demonstrates an example of sequence image normalization output. Normalized images thus allows to captures entire viral population structure using a single data representation independent of the number of sequences and with minimum loss of existing data or introduction of artificial data.

**Figure 1:**
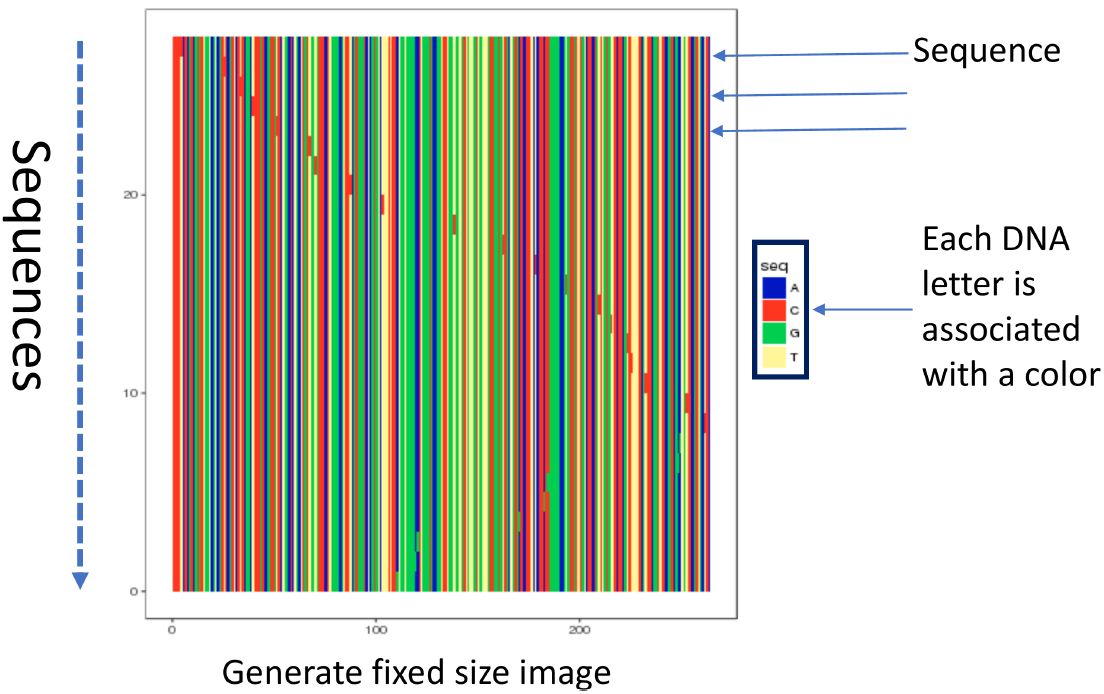
Sequence Image normalization of a fasta file

Raw pixel data of generated images are used as features to train machine learning models for the consecutive analysis, as demonstrated by Fig. 2. The number of features depends on the image resolution: each image of the resolution *x* × *y* corresponds to *x* × *y* × 3 feature vector, with each pixel having 3 RGB components. In our experiments, sequencing datasets by default has been converted into images with resolution of 480 × 480.

**Figure 2:**
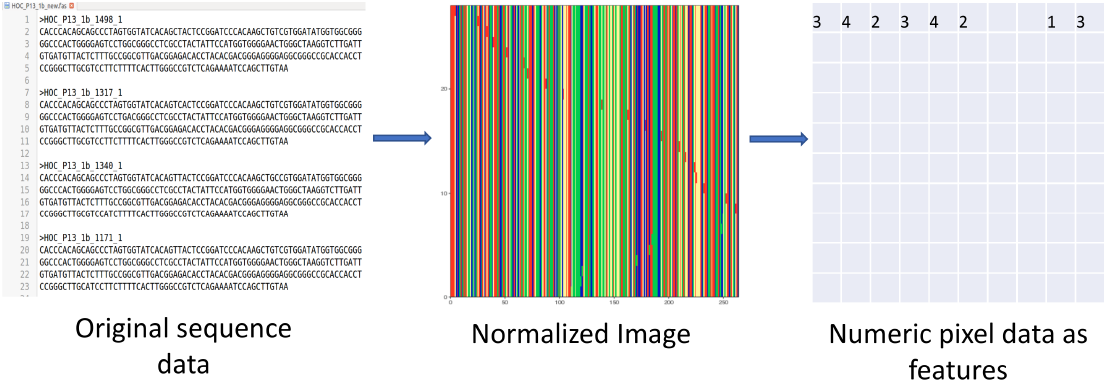
Sequence Image normalization of a fasta file

### Classification of intra-host viral populations

Identification of an HCV infection stage is considered as a binary classification problem. Images corresponding to intra-host viral populations of samples described above has been labeled based on the type of the infection as recent or chronic. This labeled data has been used to train the following machine learning classification models: Stocastic Gradient Descent (SGD), Decision Tree, Gaussian Naive Bayes(Gaussian NB), Linear Support Vector Machine (Linear SVM), Random Forest and *k*-Nearest Neighbours(*k*NN). We used models’ implementations from python `scikit-learn` library [25]. Different SVM kernels have been explored of which SVM with linear kernel produced the best results. In linear SVM model, there is a regularization parameter *c* which helps in generalizing the model by controlling testing and training errors. In this model, grid search is performed on *c* values in the range [−2, 20]. For k-NN models, we selected the best model among the models with euclidean and manhattan metrics and with *k* from the range [3, 20]. For random forest, the best model has been chosen by performing grid search on the number of trees in the range [10, 100].

Trained classifiers has been validated based on their accuracy, area under the curve (AUC), precision, and recall. Accuracy (Acc) is defined as the proportion of test cases correctly classified as either recent or chronic. Precision (Prec) measures the fraction of the correctly classified populations within each predicted infection class, while recall (Rec) measures the fraction of the true recent or chronic populations that are correctly predicted as such. Validation has been performed via stratified 10-fold cross validation. Specifically, in addition to the standard 10-fold cross-validation, we employ “leave-one-outbreak-out” cross-validation and random undersampling methods to balance the datasets. In our current data, some of the samples come from the same HCV outbreak. Such samples are close to each other by their nucleotide composition, thus their presence may lead to over-fitting of any particular method. In “leave-one-outbreak-out” cross-validation, data from each of these outbreaks was used in the validation set, while other samples are used in the training sets. Random undersampling has been performed to balance the difference in sizes of datasets of recent and chronic hosts. In this method, chronic dataset size is reduced by random subsampling to match the acute dataset size.

### Clustering of intra-host viral populations

We detect outbreak by clustering images representing intra-host viral populations. For this purpose, we employed agglomerative hierarchical clustering, *k*-means clustering and mini-batch *k*-means clustering. As before, we used models’ implementations from python `scikit-learn` library [25]. Several distance measures have been employed, including euclidean, manhattan and cosine metrics. Hierarchical clustering has been executed using complete, average and ward linkage approaches.

Normalized Mutual Information (NMI) [33], homogeneity [27] and completeness [27] scores as used as metrics to analyze the clustering performance. These measures evaluated the assigned cluster labels after clustering compared to the actual cluster class label of each intra-host viral population. Homogeneity score measures if the all members of a cluster actually belong to one cluster class label, while the completeness scores measure if all the members of an actual cluster class label are grouped into the same cluster. NMI measures the mutual information shared between the individuals in the clusters. All these measures range from 0 to 1 and the values closer to 1 refer to better clustering efficiency. To evaluate the effectiveness of the normalization method in detecting relatedness between any pair of samples, we compute AUROC (Area under ROC curve) is computed (as done in [13]). Viral populations taken from the same outbreak are considered as genetically related, otherwise as unrelated. There are 55945 pairs of samples, and 479 of them are related. After computing the distances between each pair of samples, all the pairs crossing a threshold value are considered as related. To compute AUROC curve, false-postive rate (FPR) and true-postive rate(TPR) are measured by modifying the threshold starting from the best threshold value where there are no false positives.

## Results

### Classification of infection stages

Stratified 10-fold cross validation has been initially performed to analyze the performance of several classification methods trained using the normalized image data. On Fig. 3 accuracies and AUC of the best models for each of the methods are shown using box plots, with the average metrics being emphasized by the red line. Linear SVM demonstrated superior performance compared to all other models, with an average accuracy of 97.545% and low accuracy variance. Other models with the exception of Gaussian NB have accuracies higher than 85%, thus exceeding accuracies of existing methods which are primarily based on feature extraction methods (see Comparison with previous methods subsection). Accuracy metric alone cannot justify the performance of the model as it needs to also achieve higher precision and recall metrics for each infection type. Fig. 4a - 4d demonstrate the precision and recall metrics for chronic and recent samples separately. As before, linear SVM achieved the best performance over all other models with an average precision and recall values of 98.11% and 98.45% for chronic populations and 96.52% and 95.36% for recent populations, respectively. This model also has low variance across the values obtained from all the folds. Noticeably, other models with the exception of Gaussian NB also achieve more than 80% values for these metrics.

**Figure 3:**
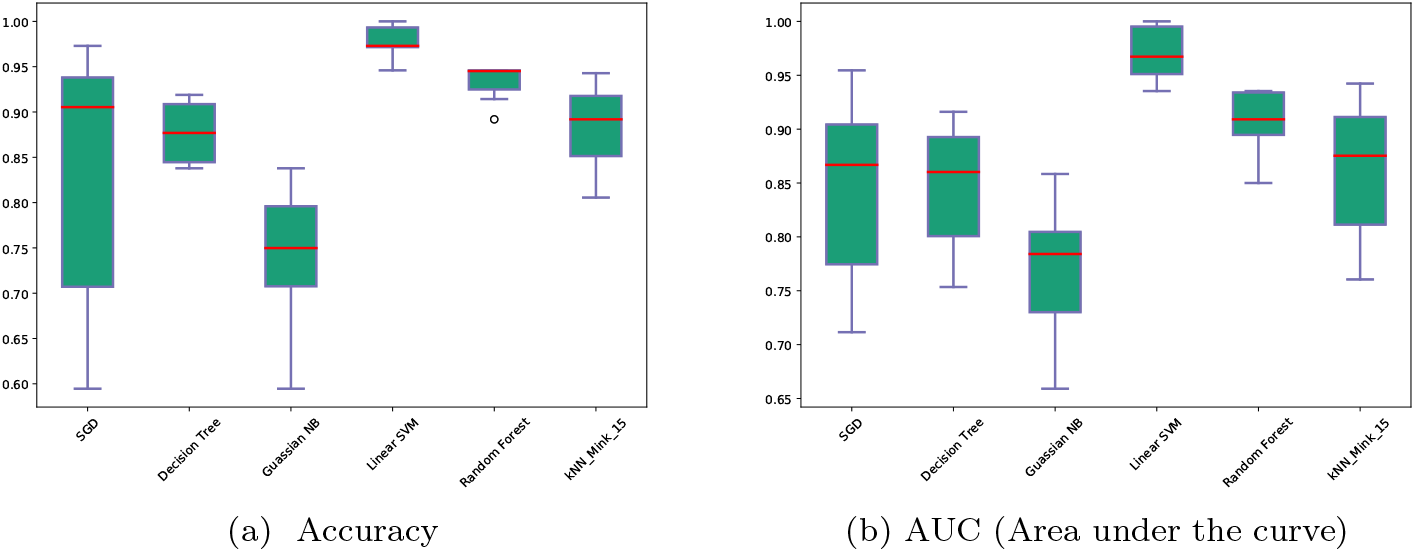
Accuracy and AUC (Area under the curve) comparisons of several simple classification methods after training them based on the normalized image data.

**Figure 4:**
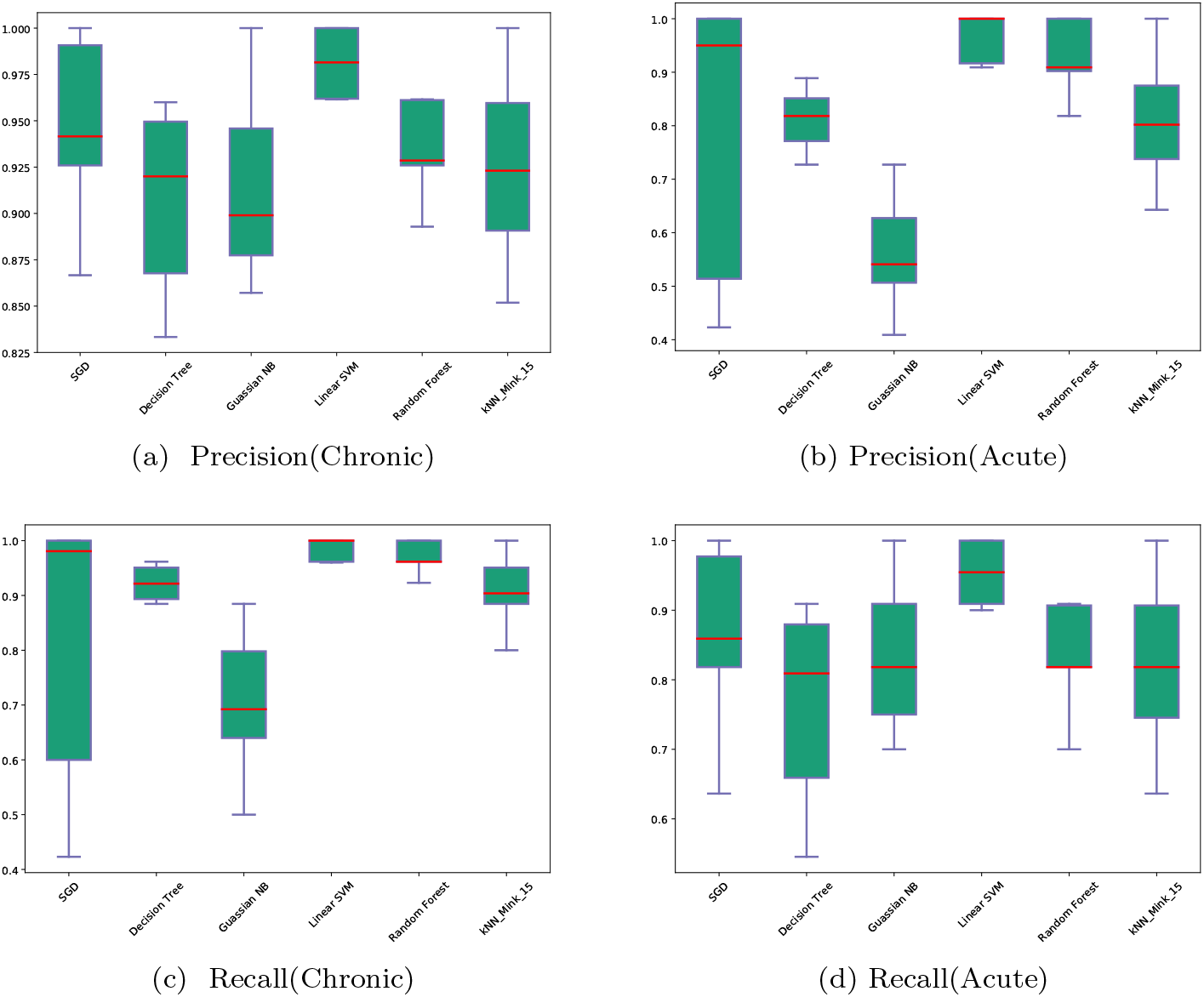
Precision and Recall comparisons of several simple classification methods after training them based on the image data generated after our sequence image preprocessing method.

Linear SVM model has been analyzed further with leave-one-outbreak-out and random undersampling validation combined with 10-fold cross-validation. Table 1 shows the results of these methods compared to the standard 10-fold cross validation on the whole dataset. The classification accuracy remains stable under the additional sampling methods.

**Table 1:** Performance metrics of Linear SVM classifier assessed by standard 10-fold cross validation, leave-one-outbreak-out validation and random undersampling methods

| Sampling Methods | Accuracy | Precision-Chronic | Precision-Acute | Recall-Chronic | Recall-Acute | AUC |
| --- | --- | --- | --- | --- | --- | --- |
| Standard 10-fold cross-validation | 97.545% | 98.105% | 96.515% | 98.446 % | 95.364% | 96.905% |
| Leave-one-outbreak-out | 96.075% | 97.004% | 91.0% | 98.446 % | 83.5% | 90.973% |
| Random undersampling | 95.164% | 96.328% | 94.661% | 94.155 % | 96.173% | 95.164% |

### Detection of transmission clusters

The results of *k*-means, mini-batch *k*-means and hierarchical clustering models are shown in Table 2. In our experiments, agglomerative hierarchical clustering with ward linkage and euclidean distance between images demonstrated the best performance. Furthermore, we evaluated the accuracy of detection of epidemiologically related pairs of populations. Two populations are considered to be related, if the distance between corresponding images is below a specified threshold. ROC curves for the accuracy of detection of epidemiologically related pairs for different distance measures and thresholds are shown on Fig. 6. All distance measures expressed consistent performance, with AUC exceeding 0.99 for all of them.

**Table 2:** Performance metrics of various clustering methods

| Clustering Method | NMI | homogeneity | completeness |
| --- | --- | --- | --- |
| *k*-means | 0.986 | 0.994 | 0.978 |
| Mini-batch *k*-means | 0.985 | 0.992 | 0.978 |
| Hierarchical | 0.987 | 0.994 | 0.979 |

### Effect of image resolution

All the experimental results discussed above have been obtained using the default image resolution 480 × 480. We analyzed the impact of image resolution on classification and clustering performance. Resolution values have been varied from 50 × 50 to 550 × 550 with step size of 50. Fig. 5a shows the performance metrics of stratified 10-fold cross validation using LinearSVM model for detecting stage of HCV infections based for different image resolutions. Highest accuracy is achieved at the resolution 450×450, although the accuracy mostly saturates approximately after the resolution 300 × 300. Similar performance has been observed for agglomerative hierarchical clustering (Fig. 5b).

**Figure 5:**
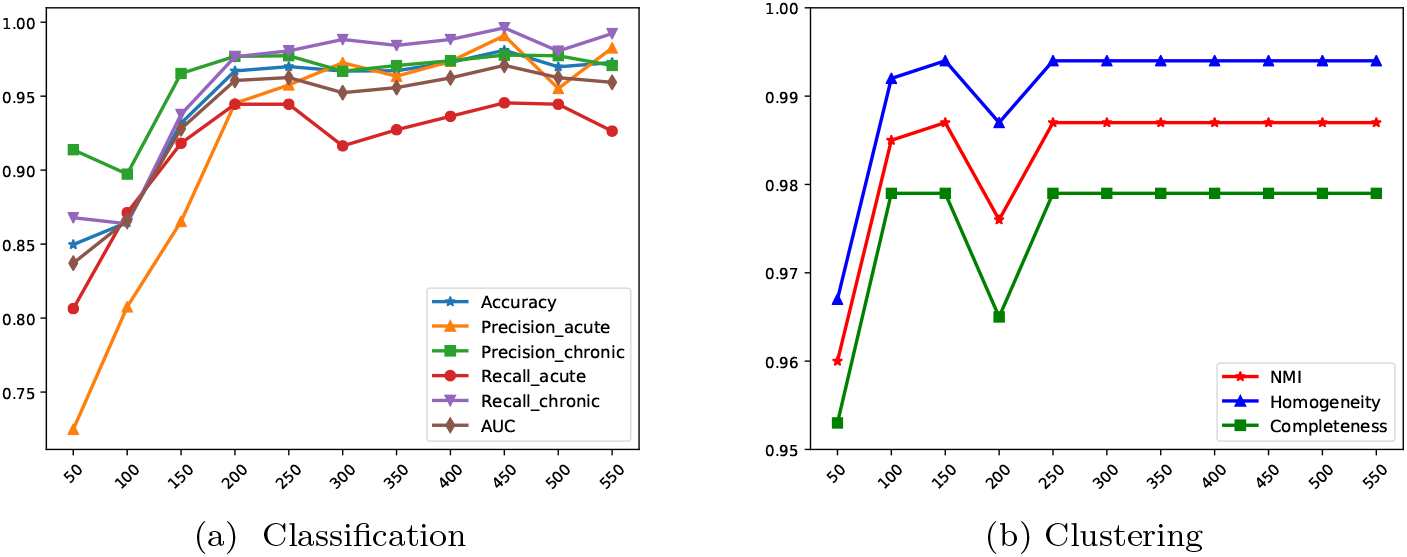
Performance metrics (Y-axis) of classification and clustering methods based on different image resolutions(X-axis).

**Figure 6:**
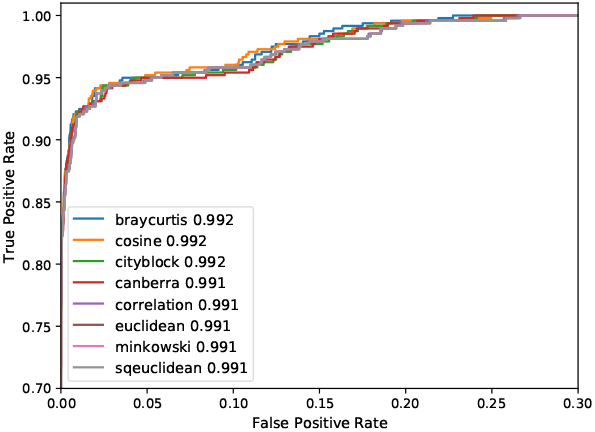
Performance of AUROC in detection of epidemiologically related pairs of populations with different distance metrics.

### Comparison with previous methods

Previously published model [24] classifies the stages of HCV infection using one of the following 3 parameters: variant frequencies entropy, average position-wise nucleotide entropy and the average distance from viral variants to the most frequent variant of the population. On our data, AUC for these parameters was equal to ~ 81%, ~ 66% and ~ 78%, respectively, while the proposed classifier based on image normalization yielded ~ 96.9% AUC.

We also compared the proposed method for the inference of genetic relatedness between different HCV samples with the two methods VOICE and ReD proposed in [13]. Image clustering method achieves sensitivity of 98.181% and AUROC of 99.2% which is similar to the VOICE algorithm and higher than ReD algorithms.

## Conclusion

In this study, we propose a novel method for generation of fixed set of features representing heterogeneous viral populations, which is widely applicable for various classification and clustering tasks addressed by machine learning. The method converts sequencing data into fixed-size images thus addressing several issues associated with comparison of viral populations by machine learning methods. The simplicity of the sequence image normalization method makes it a robust approach for converting genomic data into numerical data. The image data also helps in visualization of the original genomic data. Experimental results demonstrate that the preprocessing method converting sequencing data into images can be successfully applied to different problems from the domain of molecular epidemiology and molecular surveillance of viral diseases, with simple binary classifiers and clustering techniques applied to the image data providing better or comparable accuracies than the existing models. In future work, sequence image normalization machinery can be applied to other challenging problems in viral genomics, such as detection of co-infections and prediction of drug resistance and therapy outcome.

## Acknowledgements

PS was partially supported by NIH grant 1R01EB025022 “Viral Evolution and Spread of Infectious Disease in Complex Networks: Big Data Analysis and Modeling”. A.Z. has been partially supported by NSF Grants DBI-1564899 and CCF1619110 and NIH Grant 1R01EB025022-01. PIB was supported by GSU Molecular Basis of Disease fellowship.

